# A Novel Semi-automated Proofreading and Mesh Error Detection Pipeline for Neuron Extension

**DOI:** 10.1101/2023.10.20.563359

**Authors:** Justin Joyce, Rupasri Chalavadi, Joey Chan, Sheel Tanna, Daniel Xenes, Nathanael Kuo, Victoria Rose, Jordan Matelsky, Lindsey Kitchell, Caitlyn Bishop, Patricia K. Rivlin, Marisel Villafañe-Delgado, Brock Wester

## Abstract

The immense scale and complexity of neuronal electron microscopy (EM) datasets pose significant challenges in data processing, validation, and interpretation, necessitating the development of efficient, automated, and scalable error-detection methodologies. This paper proposes a novel approach that employs mesh processing techniques to identify potential error locations near neuronal tips. Error detection at tips is a particularly important challenge since these errors usually indicate that many synapses are falsely split from their parent neuron, injuring the integrity of the connectomic reconstruction. Additionally, we draw implications and results from an implementation of this error detection in a semi-automated proofreading pipeline. Manual proofreading is a laborious, costly, and currently necessary method for identifying the errors in the machine learning based segmentation of neural tissue. This approach streamlines the process of proofreading by systematically highlighting areas likely to contain inaccuracies and guiding proofreaders towards potential continuations, accelerating the rate at which errors are corrected.

## 1. Introduction

In recent years, the field of nanoscale connectomics has advanced considerably, with significant contributions including the hemibrain volume [1] and, most recently, the volumes collected under the Intelligence Advanced Research Projects Activity (IARPA) Machine Intelligence from Cortical Networks (MICrONS) program [2, 3, 4, 5]. Mapping brain structure and function at the neuron-synapse resolution is a key step to building our understanding of the brain. Recent modeling work, for example, has already simulated the fly at single cell resolution yielding insights into the innner workings of the optic lobe, gustatory circuit, grooming behavior, and head direction [6, 7, 8]. These efforts have generated high-resolution electron-microscopy (EM) datasets on the order of petabytes, and as ambitions are set on extracting the connectome of larger organisms like the mouse, tools need to be created to increase the throughput of error correction mechanisms. Despite developments in state-of-the-art computer vision segmentation algorithms, the resulting connectomes are imperfect and still require human expert intervention.

Several tools have been proposed to address the challenges in proofreading large-scale nanoscale connectomics datasets [9, 10, 11, 12, 13, 14, 15]. Errors in the segmentation of these datasets are often characterized as either “false splits” where two labeled regions should both have the same label or “false merges” where two regions of a single label should be separated. We refer to the correction of false splits along a neurite as “extension.” Current approaches to extension include RoboEM, a recent approach to “fly” down neurites and add extensions which works well with well aligned datasets to autonomously extend neurites [16, 17, 18]. The current work seeks to hone the process of identifying error locations cheaply to further orient both proofreaders and automated methods. For neural network based reconstruction methods, error locations generated by the current work could augment datasets as these locations often occur at the sites of image artifacts and rare cellular structures. NEURD is a Python software package designed to facilitate the analysis of EM reconstructions of neural circuits by transforming 3D neuronal meshes into extensively-annotated graph representations, providing pre-computed features that capture neural morphology and connectivity at a coarse level [19]. NEURD has demonstrated its utility in automated proofreading, proving highly effective at correcting merge errors using heuristic rules, but it does not address extension.

In this paper we propose a computationally efficient, CPU-driven, semi-automated proofreading method capable of operating on large scale connectomics datasets. We perform graph algorithms on the computed meshes of neurons of interest in order to identify regions associated with three primary mesh properties indicative of potential errors. First, we assessed vertex defects, defined as 2*π* minus the sum of the angles of every face that includes that particular vertex. This property generally locates areas that are disturbed by irregular tissue, for example an endoplasmic reticulum or an image artifact. Next, large facets can suggest segmentation errors, as dendritic and axonal surfaces at the nanoscale level typically possess fine, intricate geometries. Facets are large connected regions with the same normal vector which are generally areas where the segmentation is falsely split in the anisotropic depth dimension. Lastly, we developed a criterion we refer to as ‘encapsulated meshes,’ which are submeshes that exist entirely within the parent neuron mesh. These encapsulated meshes highlight regions where the segmentation algorithm may have missed an area that is completely enveloped by the neuron’s structure. By identifying these properties, we aim to effectively pinpoint regions that may contain errors and thereby guide proofreaders towards these potential locations to extend axons and dendrites to enhance the overall accuracy of the neural network reconstruction. The accuracy of the error detection mechanism will be demonstrated against an evaluation dataset of 15 neurons which were thoroughly extended by an expert proofreader. These measures, alongside the skeletonization of the meshes [20], generate a set of heuristics to point proofreaders towards potential errors, accelerating their error correction rate .

We will then draw implications for how to include these and other similar approaches into a semi-automated pipeline with proofreaders. Our proofreaders utilized NeuVue, a robust software platform designed for large-scale proofreading of machine segmentation and neural circuit reconstruction in high-resolution electron microscopy connectomics datasets. Its backend queuing service enhances proofreader throughput by organizing tasks into distinct, purpose-driven categories and streamlining actions into simple, atomic operations [21]. This platform enabled us to investigate the rate at which proofreaders generated edits to the segmentation both with and without the semiautomated suggestions.

## 2. Materials and Methods

### 2.1. Data and Management

This work utilizes an open-source transmission EM dataset of the mouse visual cortex consisting of 75,000 neurons and over 520 million synapses generated by the MICrONs Consortium [5]. The data was aligned and then segmented via convolutional neural networks [22]. After segmentation, each neuron can be analyzed as a mesh which translates dense 3D image matrices into a graph representation of the surface of the neuron. The meshes are processed according to the protocol detailed in [19]. The location of each soma was queried from the results of a segmentation stored in CAVEclient, a Python interface to the Connectome Annotation Versioning Engine (CAVE) APIs [16]. CAVE was also the service used to download neuron meshes and retrieve synapse counts [5].

Additional pre-processing was performed on the meshes to simplify future computation and find other classes of errors. Graph algorithms on the meshes were conducted with the help of the Trimesh Python package [23]. A connected component analysis was first conducted to extract the largest component of the graph. This discarded fragments of the mesh that result from manual merging and discontinuities in the segmentation. Some errors could be detected at this stage by determining if the vertices of each smaller mesh lie completely within the largest mesh. These fragments represent local discolorations, darkly stained organelles, and other regions which are incorrectly split from the parent’s mesh. These types of ‘encapsulated’ errors were frequent throughout the evaluation dataset, though they rarely contain synapses. The largest mesh component is then skeletonized using the TEASAR-based method utilized by the Meshparty software package [20]. The skeleton is preferred to be as course as possible in order to not include thin spines or axonal segments. Endpoints of this skeleton are calculated by selecting vertices with a degree of one. Endpoints within 3,000 nm of each other are combined to accommodate areas with many close branches, for example a false merged glial cell.

### 2.2. Defect Error Detection

Mesh defects are found in order to locate areas which are locally rough or jagged. These regions are usually not smooth due to local discoloration, dark organelles, or image artifacts which, in addition to disturbing the mesh, also often cause a false split. By searching for these jagged regions close to skeletonization endpoints, locations of possible extension can be discovered. Intuitively, vertex defects provide a measure of the “curvature” or “bending” at a vertex in a mesh surface. A vertex defect is the amount by which the sum of angles around the vertex deviates from 2*π* (the sum in a flat plane), hence representing how much the local geometry around that vertex deviates from flatness. This intuition derives from the Gauss–Bonnet theorem of geodesic triangles.

For every vertex *v*, the defect score is calculated as:

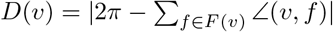

where D(v) represents the vertex defect score at vertex v. F(v) is the sum over all faces f that include the vertex v. ∠(v, f) represents the angle at vertex v in face f. However, it is difficult to find local regions of jaggedness based on each individual vertex. To “smooth” local subregions, the defect scores are transmitted through the mesh’s node adjacency matrix through two rounds of matrix multiplication and averaging.

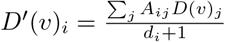

where *D*^′^ is the smoothed defect score vector, *A* is the graph’s adjacency matrix, where *A*_*ij*_ = 1 if an edge exists between vertex *i* and *j*, and *A*_*ij*_ = 0 otherwise. *d*_*i*_ is the degree of vertex *i*. This operation is performed twice to distribute defect scores to a local subregion of the graph. After this, all graph vertices with a defect score of less than 0.75 were removed. Intuitively, when this threshold is too low, many regions that do not contain errors are included, and if the threshold is too high, true error locations are omitted. After thresholding, onnly vertices surrounding high probability error locations are considered. Then, connected components are gathered on the remaining graph, and connected components with fewer than 20 vertices were discarded as they were too small to represent false splits.

The mean position is taken for the remaining components. Then, the path length along the mesh is calculated from each defect region to each skeletonization tip. Finally, it was the case that this process yielded many components which were made of long linear ridges or corners on the mesh. To ensure the regions were roughly circular or elliptical, a principal component analysis (PCA) of the coordinates was performed [24]. If the variance explained by the first component is sufficiently low, it implies at least two dimensions are needed to represent the points, meaning the points are spread out along more than one dimension. In general, large regions close to a tip are more likely to actually contain an error to be corrected. A final ranking score is then calculated for each defect subregion as:

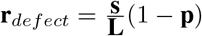

where **r** is the vector of ranking scores, **s** is the vector of the areas of each defect subregion, **L** is the vector of the path lengths to the closest skeletonization endpoint, and **p** is the vector of variances explained by the first principal components. These ranks can be used to impose a threshold on the defect error locations. Errors could also be located by removing the ranking factor **L**, these regions were often disturbed areas of the mesh where many orphaned segments could be merged with the parent mesh, but these segments rarely contained synapses.

### 2.3. Facet Error Detection

Errors in anisotropic image volumes disproportionately occur along the coarser resolution anisotropic axis. In the MICrONs datasets, this axis is the depth of the brain. Slices of tissue can be blacked out, alignment issues may exist between slices, and other types of shearing or local nonlinearities can cause these errors. In order to quickly find them, the normal vector of each face can be calculated. Then, faces that do not have a normal vector pointing exactly along the anisotropic axis can be masked out, and a connected component analysis can be performed on the remaining faces.

Once again, larger area facets are more likely to be true error locations. A ranking score again is calculated as:

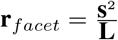

This time, the area is squared since it seemed to be more important that the facet be large than to be very close a skeletonization tip.

### 2.4. Evaluation

The method was evaluated both in the forward and reverse direction. There is a choice to make in an extension pipeline whether to start from the soma-containing neuron segments and extend outwards or to start from ‘orphan’ segments which do not have a soma. In the forward direction, the algorithm is performed as discussed above. The target neuron is preprocessed and skeletonized, and the errors are located. For the reverse direction, because they are smaller and therefore fewer possible error locations, candidate error locations were not judged based on size. Instead, errors were ranked solely based on their distance to skeleton endpoints. Similarly, we removed the ranking step from error location detection, focusing solely on filtering based on distance to skeleton endpoints and PCA.

To deploy tasks for the expert proofreaders, the system was Dockerized and given a list of neurons of interest for forward extension [25]. Tasks were queued using the Neuvue proofreading platform [21]. Before these semiautomated tasks, the expert proofreaders worked without algorithmic support. One set of tasks involved entire neurons assigned to proofreaders who were to ‘clean’ each neuron exhaustively, trying to remove as many false splits and false merges as possible. This semi-automated pipeline instead assigned each endpoint as its own task. In this framework, proofreaders received an endpoint, selected as many falsely split segments as was obvious, and then received a new endpoint.

## 3. Results

The method performed well against the expert-proofreader created dataset (Figure 2). Precision is greater than 0.8 for 12/15 of the neurons. The algorithm consistently captures a confident subset of the error locations, particularly the facet errors. Including the defect errors increases recall to cover error types that the facet detection method could not, but each defect error is somewhat less confident. Also of note, the defect error detection will sometimes find false merges, whereas the facet error detection will only detect false splits. It should be noted that while the creation of the evaluation set was thorough, the sheer size of the data means that even an expert has difficulty exhaustively cleaning neurons. There were several instances when checking the failure cases of the algorithm where the algorithm uncovered a false split that was not initially found. The creation of an exhaustive error correction training set would be a good resource for the creation of future machine learning and heuristic-based error detection tools.

**Figure 1:**
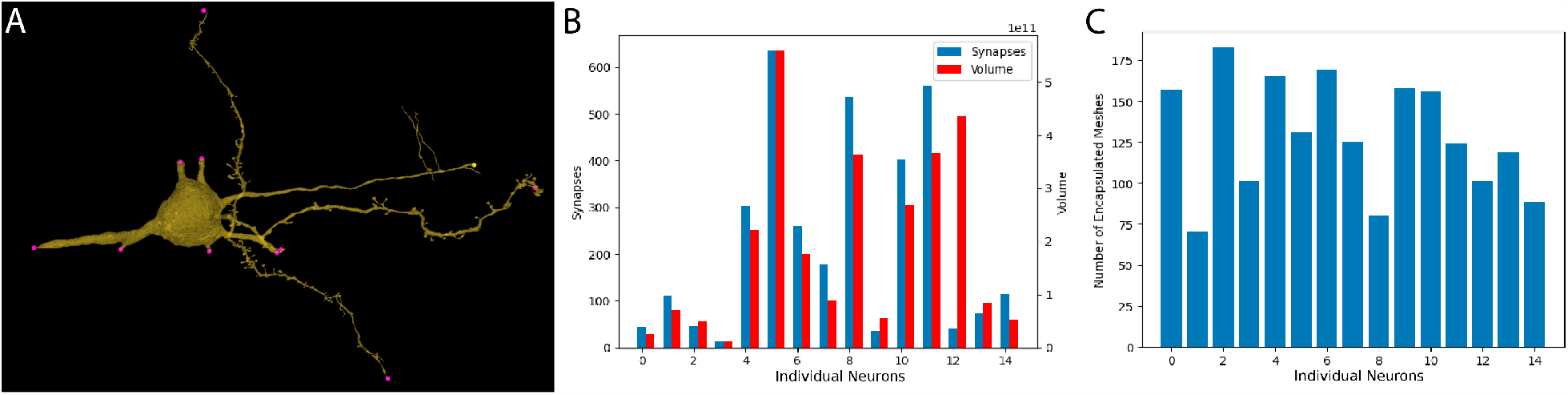
Errors in 15 expert-labelled neurons. A: A representative example of a poorly-reconstructed neuron mesh. Pink points indicate the tips of dendrites, error or not. Yellow points indicate axonal tips. B: Number of synapses and 3D volume missing from each neuron. This is the number of synapses and amount of volume located in all of the true extensions, as labeled by an expert proofreader. C: The number of ‘encapsulated’ errors within each of the expert-labeled neurons as detailed in section 2.1

**Figure 2:**
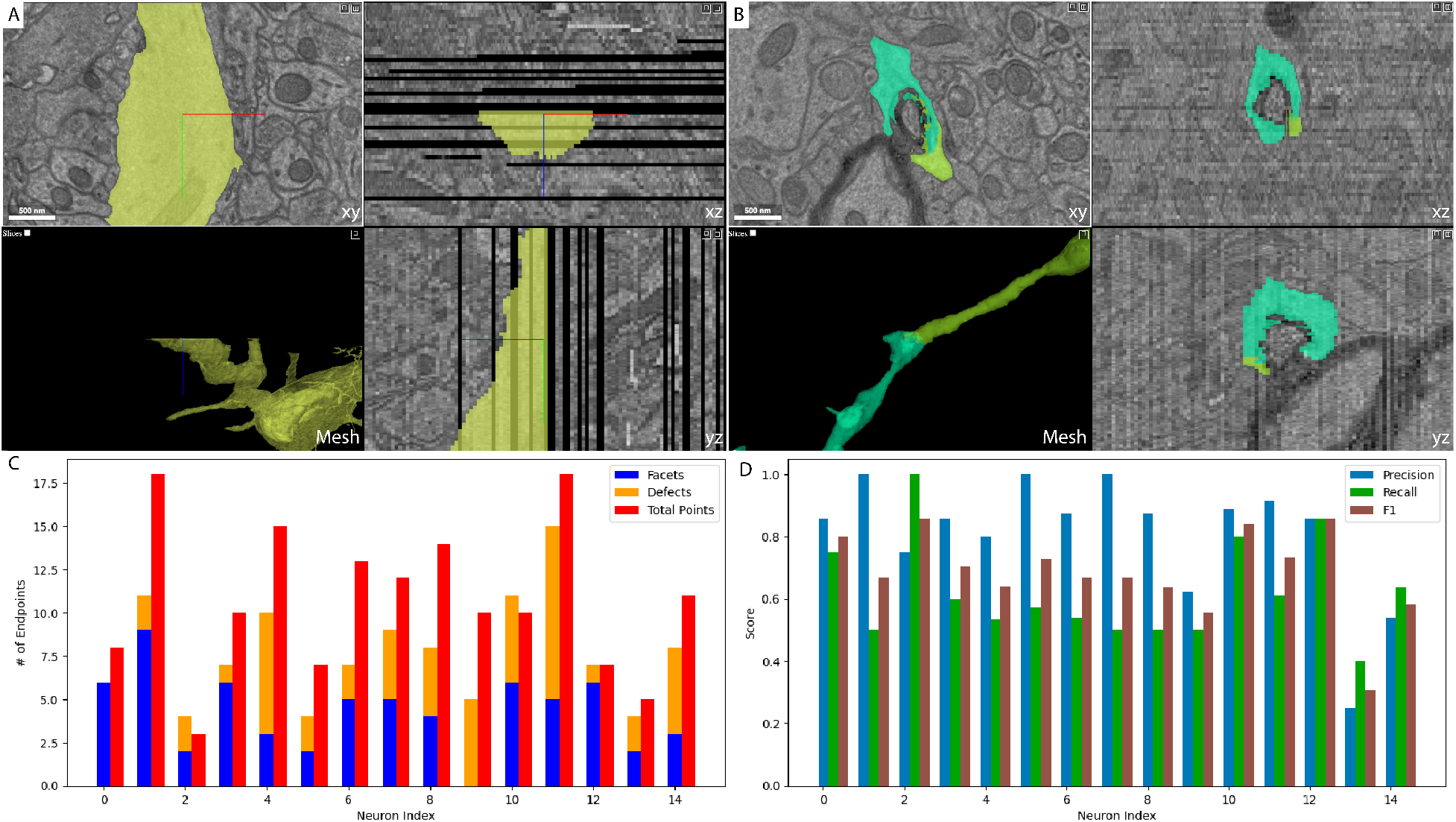
Performance of the error detector against the evaluation dataset. A: A representative example of a success case for facet detection. In the xz plane, the dropped slices that cause the split are visible. B: A representative example of a success case for defect detection. A darkly stained body interrupts the process. C: Total number of suggestions versus the total number of error locations for each neuron. D: Precision, recall, and F1 scores of the facet and defect errors together, with double counts removed.

The expert proofreaders worked notably faster when given algorithmic help (Figure 3). Their average edit rate when given no help was 0.84 edits/minute compared to 1.32 edits per minute when they were oriented towards endpoints of interest. One point of note is that because the semi-automated tasks were offered one at a time, loading time between tasks was a significant part of the time spent working. Future software improvements combined with semi-automated task generation will yield even greater task efficiency from valuable proofreader time. The expert proofreaders generated 4806 extension edits across 477 nucleus IDs, resulting in an average of 10.08 edits per neuron.

**Figure 3:**
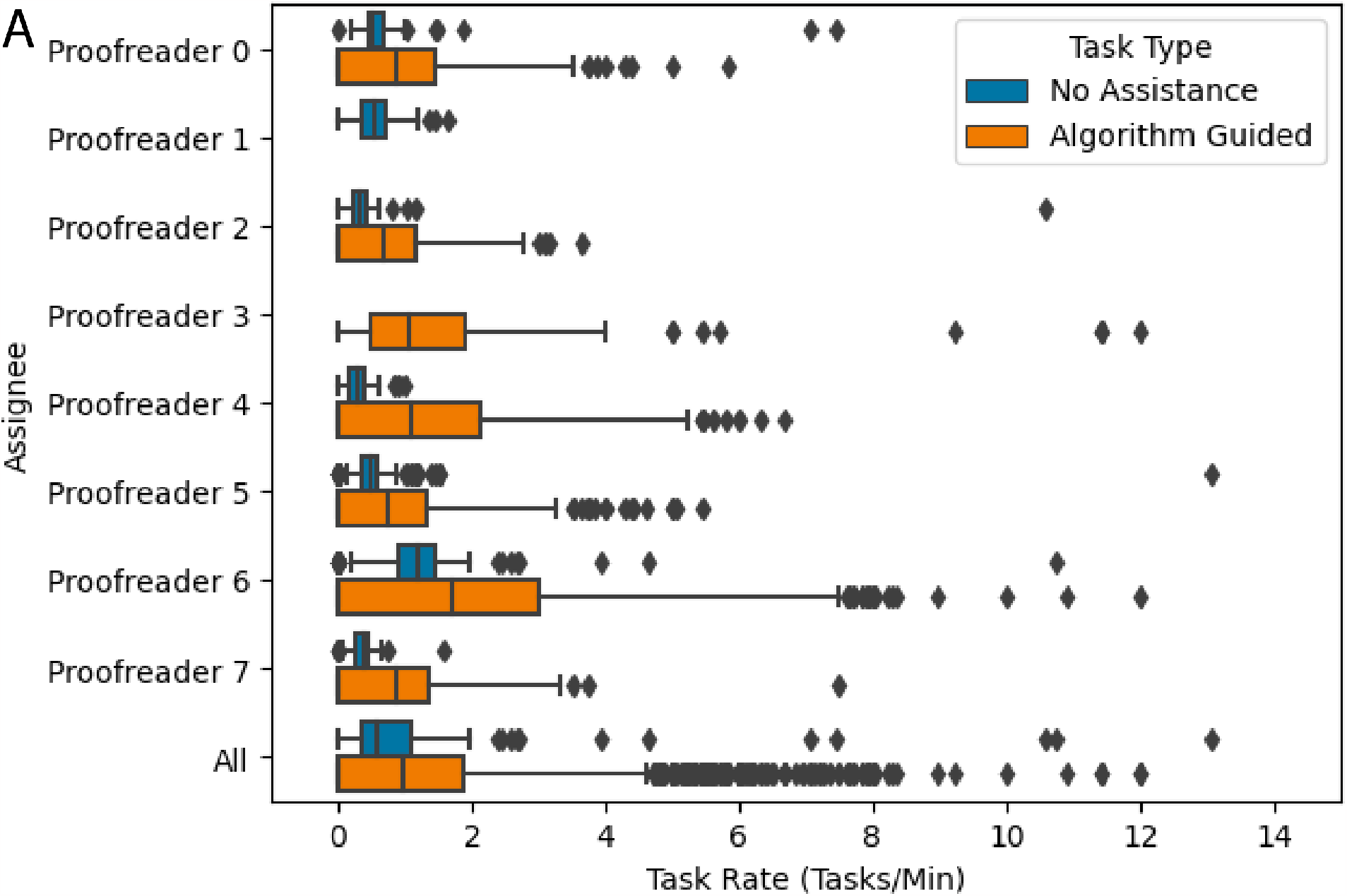
Comparison of the time each expert proofreader took on semi-automated vs manual tasks. Every proofreader who completed both task types performed faster when given automated tasks, and the automated tasks also had long tails in the positive direction, indicating that many ‘easy’ tasks were able to be completed almost immediately.

## 4. Discussion

In this study, we proposed and validated an automated approach to detect potential error locations in large-scale neuronal electron microscopy datasets. By combining mesh processing techniques and semi-automated proofreading methods, we were able to enhance the efficiency of error correction, significantly accelerating the process of data validation and reconstruction. Our methodology focused on identifying potential inaccuracies in three specific mesh properties: vertex defects, large facets, and encapsulated meshes. By pinpointing these areas, we could guide proofreaders towards potential error locations to enhance the overall accuracy of the neural network reconstruction.

The accuracy and reliability of our algorithm are affirmed by its high precision in comparison with a dataset thoroughly proofread by an expert. However, the overall recall and precision metrics suggest that there is still room for improvement in our detection methodologies. Future research should consider additional geometric features, possibly incorporating machine learning methods to predict error locations with even higher precision.

Another significant observation is the considerable improvement in the efficiency of proofreaders when guided by our algorithmic approach. While the loading time between tasks impacted overall task efficiency, it is encouraging to note that even under these conditions, the average edit rate improved significantly when proofreaders were oriented towards potential error locations. This highlights the potential of our semi-automated method in optimizing the use of valuable proofreading resources. Future software improvements should focus on minimizing loading times and streamlining the task assignment process to further augment proofreader efficiency.

We also attempted completely automated extension, using the generated endpoints as seeds for a computer vision pipeline. This approach leveraged a unique mesh-based optical flow implementation, Sofima, from Google Brain [26]. While we saw that for many examples, the automated method would select the correct extension candidate, the pipeline struggled when there were greater than three blacked out slices in a row or large misalignments. GAN based approaches or other large machine learning strategies are likely necessary for good performance on this problem.

The importance of detecting and correcting errors in large-scale nanoscale connectomics datasets cannot be overstated, especially given the increasing scale and complexity of datasets produced by ongoing neuroscience research efforts. The implementation of an effective and efficient semi-automated proofreading pipeline could have profound implications for our understanding of neural networks, accelerating the rate at which accurate and reliable reconstructions can be made. This is especially relevant given the ever-growing scale of neuronal EM datasets, as automated error detection methods become increasingly critical in dealing with the practical limitations of manual proofreading.

## Acknowledgments

This material is based upon work supported by the Office of the Director of National Intelligence (ODNI), Intelligence Advanced Research Projects Activity (IARPA), via IARPA Contract No. 2020-20081800401-023 under the MICrONS program. The views and conclusions contained herein are those of the authors and should not be interpreted as necessarily representing the official policies or endorsements, either expressed or implied, of the ODNI, IARPA, or the U.S. Government. The U.S. Government is authorized to reproduce and distribute reprints for Governmental purposes notwithstanding any copyright annotation therein. Research reported in this publication was also supported by the National Institute of Mental Health of the National Institutes of Health under Award Number R24MH114799 and R24MH114785. The content is solely the responsibility of the authors and does not necessarily represent the official views of the National Institutes of Health. We would also like to thank the JHU Applied Physics Laboratory CIRCUIT program for allowing us to collaborate with interns and use this project to support their scientific development.

Additionally, we would like to thank Nathan Drenkow, and Erik C. Johnson for supporting previous foundational work and for valuable discussions.

